# Stability of asymmetric cell division under confinement: A deformable cell model of cytokinesis applied to C. elegans development

**DOI:** 10.1101/2022.03.17.484706

**Authors:** Maxim Cuvelier, Wim Thiels, Rob Jelier, Bart Smeets

## Abstract

Cell division during early embryogenesis has been linked to key morphogenic events such as embryo symmetry breaking and tissue patterning. It is thought that boundary conditions together with cell intrinsic cues act as a mechanical “mold”, guiding cell division to ensure these events are more robust. We present a novel computational mechanical model of cytokinesis, the final phase of cell division, to investigate how cell division is affected by mechanical and geometrical boundary conditions. The model reproduces experimentally observed furrow dynamics and predicts the volume ratio of daughter cells in asymmetric cell divisions based on the position and orientation of the mitotic spindle. We show that the orientation of confinement relative to the division axis modulates the volume ratio in asymmetric cell division and quantified the mechanical contribution of cortex mechanics, relative to the mechanical properties of the furrow ring. We apply this model to early *C. elegans* development, which proceeds within the confines of an eggshell, and simulate the formation of the three body axes via sequential (a)symmetric divisions up until the six cell stage. We demonstrate that spindle position and orientation alone can be used to predict the volume ratio of daughter cells during the cleavage phase of development. However, for compression perturbed egg geometries, the model predicts that the change in confinement alone is insufficient to explain experimentally observed differences in cell volume, inferring an unmodeled underlying spindle positioning mechanism. Finally, the model predicts that confinement stabilizes asymmetric cell divisions against bubble-instabilities, which can arise due to elevated mitotic cortical tension.

**Author summary:** A crucial morphogenic step during early embryonic development is symmetry breaking in the embryo. For *C. elegans* the formation of the three body axes can be traced back to the six cell stage, where tissue-topology is the result of symmetric and asymmetric divisions. How cell mechanical boundary conditions and cell intrinsic cues influence this process of symmetry breaking is still an open question, as currently, a quantitative mechanical description of cytokinesis in complex architectures is lacking. We developed a simple mechanical model of cell division, incorporated in an existing mechanical cortex model, to simulate cytokinesis in geometrically confined environments. Our approach was able to both capture furrow ring dynamics and predict the volume ratio of daughter cells accurately. By simulating early *C. elegans* development with different geometrical boundary conditions, we were able to trace back the origin of volume discrepancies between the experimental setups to a quantifiable shift in spindle positioning during cytokinesis. Finally, we showed how embryo confinement partially stabilizes bubble-instabilities that arise during asymmetric cell division during the early cleavage phase.

## Introduction

Early embryogenesis is characterized by a well-orchestrated sequence of cell division and differentiation events. In bilateria, an important step during this process is the establishment of the three main body axes: Anterior-Posterior, Dorso-Ventral and Left-Right [1]. In model organisms with an invariant developmental cell fate, such as the nematode *Caenorhabditis elegans*, the origin of these symmetry breaking steps has been traced back to asymmetric cell divisions, which can be biochemical and/or physical in nature [2, 3]. Although the importance of physical asymmetry, such as an asymmetric volume ratio of daughter cells, remains poorly understood, recent observations for P0 divisions in *C. elegans* have linked daughter cell size asymmetry to changes in temporal cell cycle coordination, tissue topology, cell fate, and even overall robustness of early embryogenesis [3].

To control the volume of daughter cells, cell division relies on the precise positioning of the spindle apparatus, as the future cleavage site aligns with the spindle midplane [4–6]. Accumulated discrepancies during this process can lead to polyploidic cells, which have major detrimental effects on tissue homeostasis and development [7–9]. It is thus imperative for a dividing cell to have strict spatiotemporal control of its spindle, as any shift in spindle position influences the volume ratio daughter cells and viability [10]. Multiple spindle positioning mechanisms have been identified and categorized based on the nature of the underlying cues e.g., internal vs external and biochemical vs mechanical [11]. Spindle positioning in the P0 division of *C. elegans* for example, which is a highly asymmetric both in volume and distribution of molecular species, is governed by a biochemical gradient which induces a bias in the magnitude of cortex-spindle force interactions towards the posterior pole [12]. For wild type *C. elegans*, this bias produces a net spindle division plane offset, from the cellular center-of-mass in the direction of the posterior pole. This offset eventually results in an asymmetric division where the anterior cell is significantly larger than the posterior cell, at a *±* 60*/*40 volume ratio. Disrupting biochemical cues, e.g. the distribution of PAR proteins, or interfering with mechanical cortex-spindle interactions can lead to symmetric P0 division [3, 13]. Interestingly, *C. elegans* embryogenesis seems robust to compression induced eggshell deformation [14]. However, in these geometrically perturbed setups, changes in cellular movement [14] and enhanced volume asymmetry in divisions that contribute to the LR-axis [15] have been observed in the early embryo.

Mechanically, the final phase of cell division, cytokinesis, is driven by a tensile furrow ring consisting of cross-linked actin filaments [16]. The dynamic cross-linking of tread-milling actin subunits in the ring is thought to drive tension build-up, resulting in furrow ingression [17]. However, the specific contribution of actin-motor cross-linkers such as non-muscle myosin-II is an ongoing debate [16]. As of yet, while key biochemical mechanisms that regulate cytokinesis have been identified [2, 18–21], a full spatiotemporal mechanical description of cytokinesis at cell scale is still lacking. The past decade has however seen significant progress in the development of mathematical and computational models that describe the physics of cytokinesis. However, most fail to account for the geometrical and mechanical constraints imposed by the surrounding tissue architecture and the embryonic boundary. Furthermore, few mechanical models exist that are able to represent the physics of asymmetric cell division [22, 23]. Yet, especially in early embryonic development, it is thought that the coupling between the asymmetry of cell divisions and the geometry and connectivity of the surrounding tissue stabilizes robust tissue patterning [24–28]. At long timescales, the model of a cell as a fluid droplet with surface tension and adhesive bonds provides a minimal description of the mechanical response of a cell. It can be tuned with a small set of experimentally accessible parameters, such as the interfacial tension and the effective cortical tension [29, 30]. However, droplets under tension are inherently susceptible to “bubble-instabilities”, where regions with locally elevated curvature, such as the small daughter half in an asymmetric division, tend to collapse in favor of low curvature regions due to differences in local pressure. Interestingly, such instabilities have been observed *in vitro* in HeLa cells and fibroblasts [31], where they produce shape oscillations during cytokinesis.

We present a novel computational model of cytokinesis based on an existing Deformable Cell Model (DCM) [32–35]. In the DCM, the cell cortex is approximated as a viscous shell under surface tension and cell-cell interactions are takes into account through adhesion forces. A complete description of the DCM can be found in S1. For this work, we made abstraction of the underlying molecular mechanisms of furrow ring mechanics and used a coarse-grained representation where the furrow ring was approximated by a closed loop consisting of elastic subunits, which shrink as cytokinesis progresses [36], see Models section. Furrow ingression then follows as a result of elastic tension build-up in the contractile ring and local cortex deformation. Ring contraction progresses until cell abscission is initiated at the prescribed abscission radius. Following cell abscission, membrane and cortex wound healing wrap up cell division, splitting the mother cell into daughter cells [37]. *C. elegans* was used as our biological model system due to its invariant developmental path and transparent eggshell, allowing for reliable and direct volume estimates for each cell. Furthermore, significant effort has been made in mechanically characterizing *C. elegans*, permitting proper model calibration and relevant quantitative comparisons between simulation results and experimental measures.

## Results

We first validate our cell division model by comparing simulation results to experimental data and theoretical predictions found in literature [6, 17, 31, 36]. We start by investigating furrow ring dynamics, before focusing on spindle positioning and the volume of daughter cells.

### Furrow ring dynamics and cytokinesis duration

Carvalho et al. observed furrow ingression rates to be constant, until the ring reaches a threshold radius *R*_*t*_ = 3 − 3.6 μm. Beyond this threshold, the furrow ring interactions with interpolar microtubuli become non-negligible, slowing down the rate of ingression until reaching a halt at radius *R*_*f*_ = 0.5 − 1.0 μm [36]. Cell abscission and membrane fusion follow, physically separating the cytoplasm of the daughter cells [37].

To compare our simulations to the experimental work of Carvalho et al., we simulated the division of unconfined, spherical cells with different sizes and analyzed the constriction rate and cytokinesis duration, see Fig. 2. The division model parameters are summarized in Table 1. As reported in literature, cytokinesis starts with an initially constant furrow constriction velocity, see Fig. 2B, until reaching the transition radius, after which the constriction rate decreases linearly with the ring perimeter, see Fig. 2C. The total duration of cytokinesis increases for smaller cells, as the transition radius is reached earlier during cytokinesis. This scaling was also observed by Carvalho et al., which put forward an estimate for the total duration of cytokinesis as

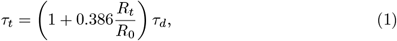

with *τ*_*d*_ as the base ring closure time and *τ*_*t*_ the expected furrow ring closure time. Model input parameter *τ*_*d*_ = 263 s was estimated based on the work of Carvalho et al. [36], see Models section.

**Table 1.** Reference table of division model parameters Parameter Symbol Value Units Deformable Cell Model.

| Parameter | Symbol | Value | Units |
| --- | --- | --- | --- |
| **Deformable Cell Model** |  |  |  |
| Cell radius | $R_0$ | 5. – 20. | $\mu\text{m}$ |
| Interphase surface tension | $\gamma_I$ | $\gamma_M/3$ | $\text{mN } \mu\text{m}^{-1}$ |
| Cytosolic bulk modulus | $K'$ | 2. | kPa |
| Effective cortex viscosity | $\eta$ | 25. | kPa s |
| Effective cortex thickness | $t$ | 300. | nm |
| **Cytokinesis Model** |  |  |  |
| Transition radius | $R_t$ | 3.6 | $\mu\text{m}$ |
| Abscission radius | $R_f$ | 0.8 | $\mu\text{m}$ |
| Mitotic surface tension | $\gamma_M$ | 1.2 | $\text{mN } \mu\text{m}^{-1}$ |
| Daughter-half bulk modulus | $K^{div}$ | 0.5 | kPa |
| Furrow ring stiffness | $k^{el}$ | 1000. | $\text{mN } \mu\text{m}^{-1}$ |

Division model parameter settings used for the cytokinesis duration and asymmetric division simulations. For a complete list of DCM parameter settings, see S2.

**Fig 1.**
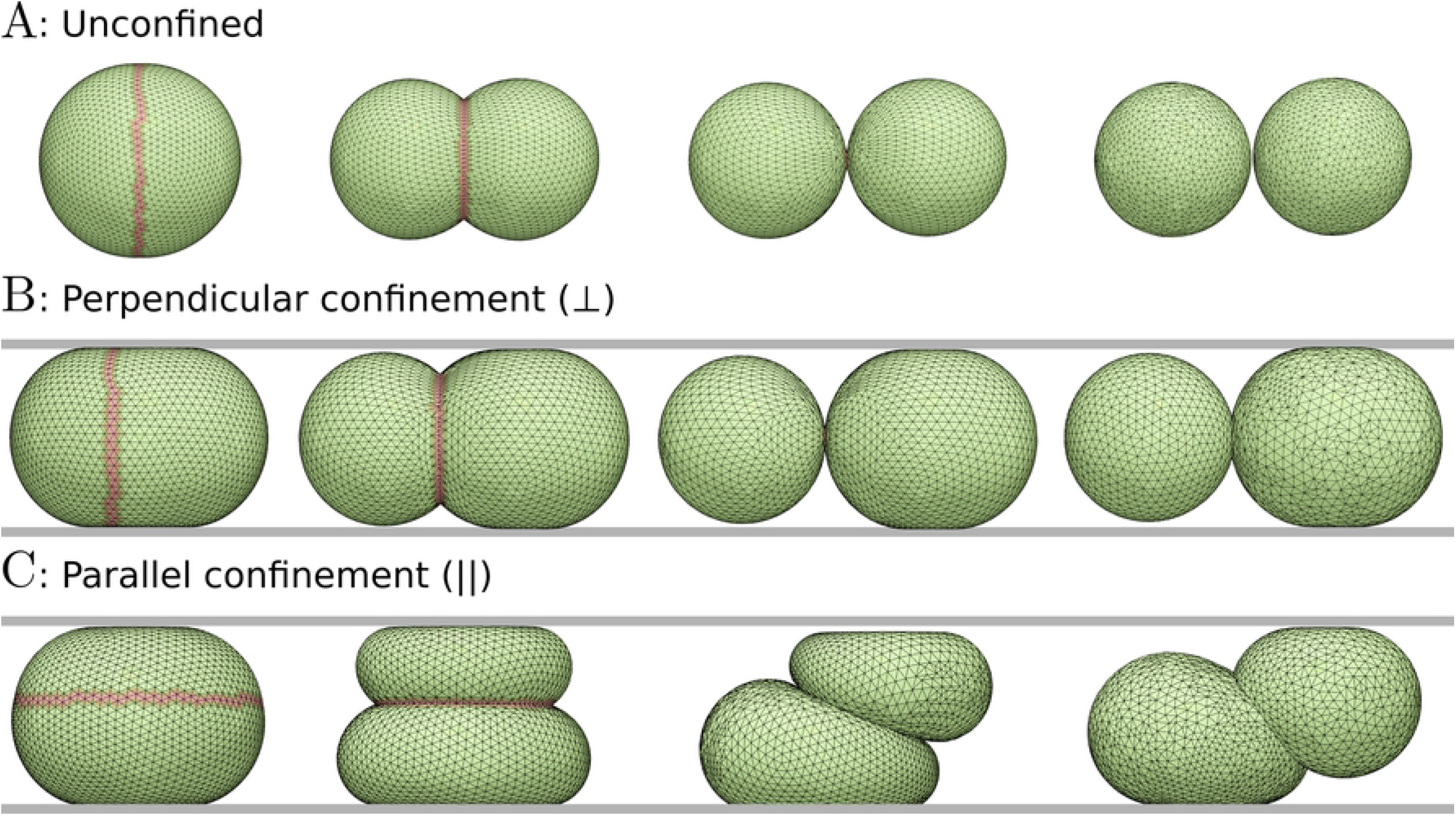
Cell shape during simulated cytokinesis. Simulation snapshots at different time points during cytokinesis. **A**: unconfined, **B**: Perpendicular (⊥) confinement and **C**: Parallel (‖) confinement. Triangles show the discretization of the cell cortex. Triangles colored in red are marked as furrow ring. In these region, tension is generated to drive furrow ingression. After abscission, the cell is split into two daughter halves and a new computational mesh is generated.

**Fig 2.**
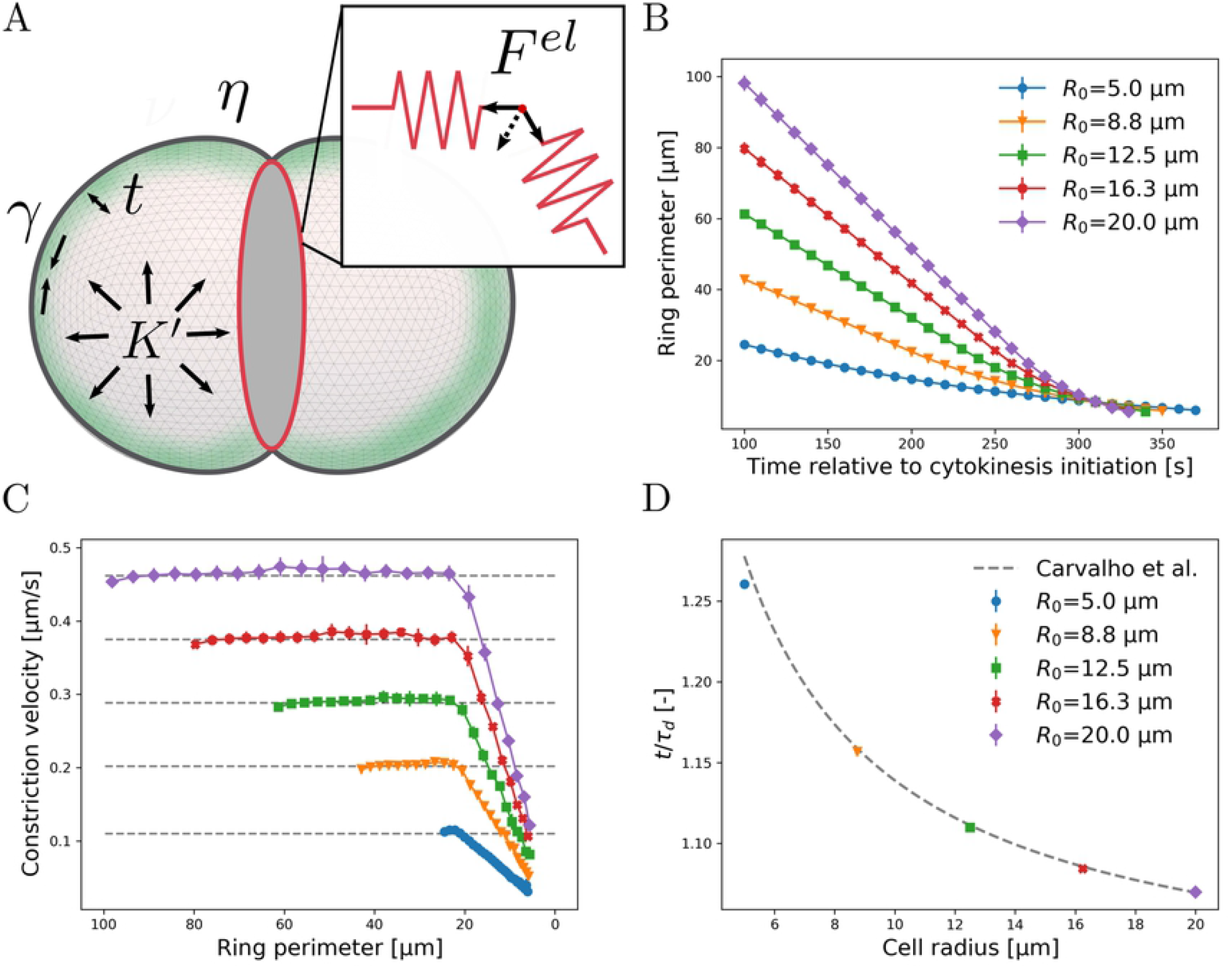
Simulated furrow constriction rate and perimeter length. **A**: Schematic representation of a dividing cell. Triangulated surface meshes are used to represent the viscous cortical shell with thickness *t*, passive mechanical properties (viscosity *η*) and active mechanics (contractility *γ* and bulk modulus *K*′) [33, 34]. Distinction is made between interphase *γ*_*I*_ and mitotic surface tension *γ*_*M*_, fixing *γ*_*I*_ = *γ*_*M*_ */*3. The furrow ring, represented in red, virtually splits the mother by generating contractile forces per sub-unit, resulting in a net inward facing force. **B**: Contractile ring perimeter in function of time during cytokinesis. Larger cells spend proportionally more time in the constant constriction regime. **C**: Constriction rate in function of ring perimeter. A constant and linear constriction regime can be identified, in line with experimental observations. **D**: Total cytokinesis duration in function of mother cell radius, together with Eq. (1), normalized with the theoretical division duration.

As our simulations were able to reproduce experimentally observed scaling of *τ*_*t*_, see Fig.2D, we conclude that our implementation of the furrow ingression model, in combination with a viscous cortical shell model [32–34], provided an adequate meso-scale description of the dynamics of cell shape during cytokinesis. We performed a sensitivity analysis, see S2, to assess the influence of the surface tension (*γ*_*M*_) and viscosity (*η*) of the cortex during mitotic rounding, relative to the stiffness of the contractile ring *k*^*el*^. In line with simulation results by Turlier et al. [22], we found that the furrow ring must be mechanically dominant, i.e., *γ*_*M*_ */k*^*el*^ ≪ 1 and (*ηR*_0_)*/*(*τ*_*d*_*k*^*el*^) ≪ 1, to be consistent with experimental observations.

### Asymmetric cell division under confinement

During cytokinesis, spindle positioning determines the volume ratio of future daughter cells [6]. To replicate this in simulations, we implemented a phenomenological description of the division plane positioning mechanism where we use cell polarization and center-of-mass, in conjunction with a spindle offset *α*, and the orientation *ϕ* of the division plane caused by active torque generation in the cell cortex, see Fig. 3A [38, 39].

**Fig 3.**
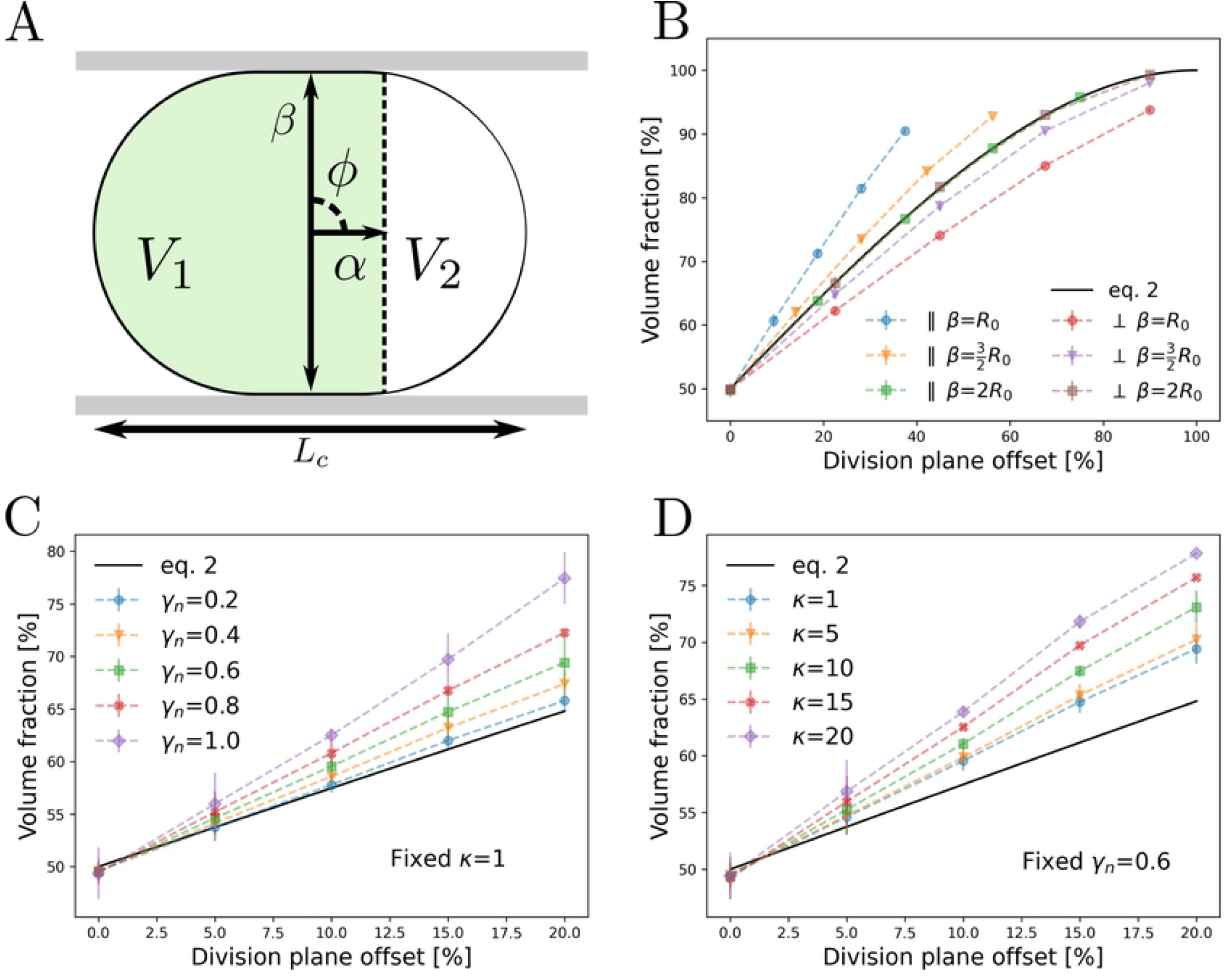
Asymmetric division under confinement. **A**: Schematic illustration of geometric parameters in the asymmetric cell division model, with characteristic length *L*_*c*_, spindle offset *α* and angle *ϕ*. For the confined simulations, two parallel plates with a gap *β* were added. **B**: Geometrically confined setup where division mechanics dominate over cortex mechanics, *γ*_*n*_ = 0.1 and *κ* = 0.5. For confinement parallel to the division plane, *ϕ* = 0, decreasing gap *β* increases daughter cell asymmetry when compared to eq. 2. Perpendicular confinement, *ϕ* = *π/*2, reverses the observed trend. **C**: Effect of mitotic surface tension on division asymmetry. Increasing surface tension destabilizes asymmetric divisions as the pressure difference over the daughter halves increases (bubble-instability). **D**: Effect of relative cortical viscosity on division asymmetry.

For spherical cells, the relative offset of the division plane, *α*_*n*_ = *α/R*_0_, with respect to the center-of-mass and daughter volume is given by

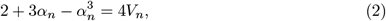

where *V*_*n*_ represents volume fraction of the largest daughter cell with respect to the mother cell. For regular geometrical shapes, similar relations can be derived. We simulated asymmetric divisions for unconfined and confined cells, verifying the agreement with eq. 2. For geometrically confined cells, a distinction was made between parallel (‖) and perpendicular (⊥) confinement, where the orientation angle *ϕ* was defined based on the angle between the division plane and confining parallel plates, see Fig. 1B,C and Fig. 3A,B. For all setups, *β* = 2*R*_0_ represents an unconfined state while for *β* = *R*_0_ the cells are compressed until the parallel plates are spaced one cell radius apart. The estimated volume ratio for unconfined simulation setups agrees well with eq. 2. For decreasing *β* in the parallel confined configuration, daughter cell asymmetry becomes increasingly sensitive to spindle offset *α*_*n*_. The opposite is observed for perpendicularly confined cells, where the volume of daughter cells are less affected by *α*_*n*_. This effect can be attributed to the scaling of the characteristic length of the cell in the direction perpendicular to the division plane, which scales as ∼ *β* for parallel confined cells and as ∼ 1*/β* for the perpendicular configuration. Finally, we assessed the effect of mitotic cortical surface tension (*γ*_*M*_) and viscosity (*η*) on the volume ratio in asymmetric cell division, see Fig. 3C,D. In the liquid droplet model of a cell, these properties drive the bubble-instability of an asymmetric division, i.e., any asymmetry will be further increased, potentially leading to complete collapse of the smaller daughter half. We postulate that a biological cell escapes this instability through a non-zero bulk rigidity of the cytosolic content, which we implemented in our model by means of a daughter-half bulk modulus *K*^*div*^. In accordance, we define the relative mitotic tension compared to *K*^*div*^ as *γ*_*n*_ := 2*γ*_*M*_ */*(*R*_0_*K*^*div*^), a measure that expresses the balance between surface tension and bulk rigidity. For increasing *γ*_*n*_, we observe increasingly asymmetric divisions, matching experimental observations [31]. The cause of the observed shift in volume ratio can be traced back to the underlying stabilizing mechanism which fails to correct discrepancies in daughter half volumes as *γ*_*n*_ increases, see Models section. Similar to *γ*_*n*_, increasing the relative contribution of cortical viscosity, *κ* := *ηR*_0_*/*(2*γ*_*M*_*τ*_*d*_), results in increasing asymmetry. In this case, however, the shift in volume ratio is caused by a change in cell shape: For high *κ*, viscous stress in the cortex is significant and influences the shape of the cortex by increasing the characteristic timescale of cortex relaxation.

### Spindle positioning during early *C. elegans* development

We apply our mechanical model for cytokinesis to asymmetric cell divisions that take place during early *C. elegans* development. We focus on the cleavage phase up to the six cell stage, as development up to this stage is thought to be predominantly the result of passive cell mechanical and cell-cell interactions, and less of active mechanical processes such as cell migration [40]. We investigated both compressed and uncompressed embryo shapes. Embryo compression occurs naturally *in-utero*, while *in-vitro* it generally occurs as a result of sample preparation for imaging, where the degree of compression depends on the mounting technique used [14]. The initial geometry of the compressed embryo was based on segmented egg shapes obtained from microscopy data [15], while the uncompressed shape was estimated by fitting an ellipsoid with the same volume [41]. The spindle offset *α* for each cell division was statistically sampled from experimental data in compressed conditions from Fickentscher and Weiss [6]. Furthermore, we assume fixed spindle skew angles *ϕ*, with values from experiments of Pimpale et al. [39]. In simulations, we assume that cell-cell adhesion is small relative to cortical tension in early *C. elegans* development [42, 43]. A full description of the computational model of *C. elegans* development is provided in the SI.

With these parameters, together with well-characterized cell division timings, we simulate the early development of *C. elegans* up to the six cell stage. The simulated development reproduces key morphogenic features of early *C. elegans* development, such as the characteristic diamond configuration in the four cell stage and dorso-ventral symmetry breaking in the six cell stage, see Fig. 4A. The volume of simulated cells were compared to volume estimates based on segmented microscopy images of compressed embryos [15], Fig. 4B. We observe that the simulated volume ratio agrees well with experimental observations. Based on this, we conclude that the proposed DCM of cytokinesis captures the dominant mechanical processes that drive asymmetric cell division well. This supports the notion that position *α* and angle *ϕ* of the mitotic spindle are highly robust predictors for volume asymmetry in the *C. elegans* cleavage phase.

**Fig 4.**
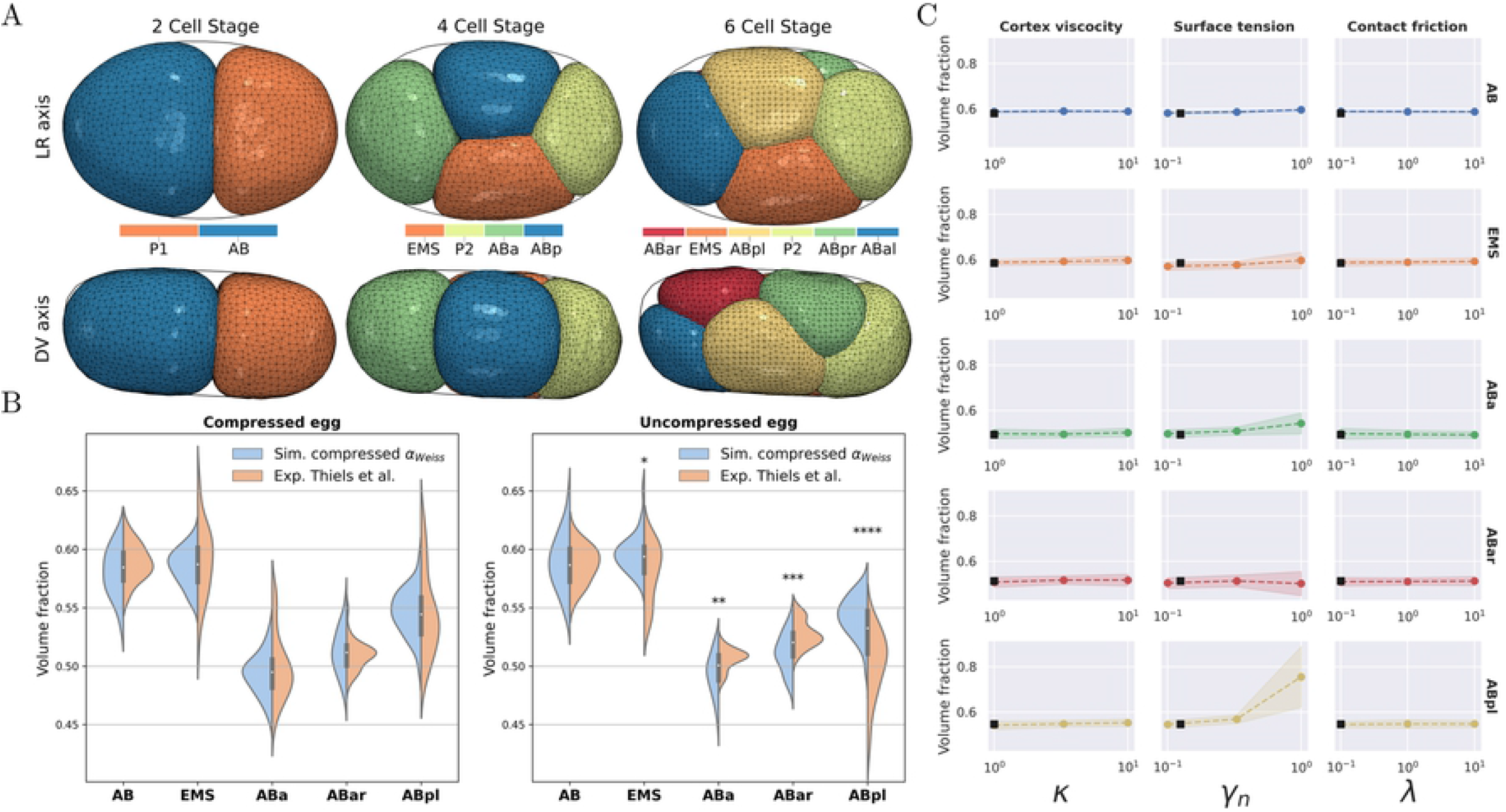
Simulated volume ratio in *C. elegans* early development. Volume of daughter cells relative to the volume of the mother cell. Only the volume ratio of the largest daughter cell is reported as the sum of larger and smaller daughter cell is always ≈ 1. **A**: Simulation snapshots for the 2-6 cell stage as seen in the LR direction (Top view) and DV direction (Side view). **B**: Simulation estimated cell volume based on *α* values reported by Weiss et al. for compressed and uncompressed eggs. For the P0 division, estimates by Gotta et al. were used [44]. Significant differences between simulation and experiment are indicated for a standard 2-sided t-test (p-values 0.05 ≤ * ≤ 0.01 ≤ ** 0.001 ≤ *** ≤ 0.0001 ≤ ****). **C**: Sensitivity analysis of daughter cell volume, for normalized cortex viscosity, surface tension and contact friction. Black squares mark the reference value of a given parameter. It should be noted that changes in cell-cell connectivity and cell positions were not quantified.

We hypothesized that the effect of confinement alone is sufficient to explain the different volume ratio observed in experiments [15]. To this end, we adopted spindle positions from experiments in confined conditions, but simulated development in an uncompressed egg geometry. However, we observed that simulation results were significantly different from experimental observations, see Fig. 4B. These results suggest that apart from cell shape changes, egg geometry also influences spindle positioning, as the shape changes alone were found to be insufficient to induce the experimentally observed differences in volume. Since spindle positions seem to be the dominant factor in determining daughter volume ratio, our model suggests that any significant change in the volume of daughter cells should be accompanied by a shift in spindle position.

Moreover, taking into account the results presented in the previous section, we expect that the sensitivity of volume ratio to the absolute spindle position depends on the characteristic length scale of the mother cell in the division direction. Hence, this sensitivity increases as development progresses and may depend on the specific geometric configuration of a cell as it initiates cytokinesis. In a complementary approach, the model was used to estimate spindle offset *α* based on experimentally measured cell volume, using a reverse fitting procedure, see S2. The results of this procedure are summarized in Table 2. This yields estimates of *α* for the compressed conditions that agree with experimental measurements [6, 44], and produces a computational prediction for the adjusted spindle offset in uncompressed conditions. In the latter case, the model predicts a mean absolute shift in *α* of *±* 300 nm. The estimated shift is statistically significant (2-sided t-test p ≤ 0.05) for the AB cell line but not for the P cell line.

**Table 2.** Estimated spindle offset *α* based on linear fitting.

| Cell | Exp. ($\mu\text{m}$) | Exp. Source | Comp. sim. ($\mu\text{m}$) | Uncomp. sim. ($\mu\text{m}$) |
| --- | --- | --- | --- | --- |
| P0 | $2.630 \pm 0.701$ | [44] | $2.883 \pm 0.157$ | $2.603 \pm 0.100$ |
| AB | $-.351 \pm 0.312$ | [6] | $-.274 \pm 0.028$ | $.077 \pm 0.051$ |
| P1 | $1.335 \pm 0.268$ | [6] | $1.398 \pm 0.114$ | $1.284 \pm 0.061$ |
| ABa | $.357 \pm 0.278$ | [6] | $.277 \pm 0.052$ | $.546 \pm 0.089$ |
| ABp | $-.617 \pm 0.255$ | [6] | $-.653 \pm 0.048$ | $-.178 \pm 0.036$ |

Estimates for $\alpha$ , using $\phi$ data from Pimpale et al. and volume estimates by Thiels et al. as input [15, 39]. A linear fitting procedure was used where $\alpha$ values were estimated via interpolation, see S2.

Finally, a sensitivity analysis was performed with respect to the influence of cortex mechanics on cell division asymmetry in the *C. elegans* embryo, thereby assessing the mechanical sensitivity of asymmetric cell divisions in confined environments, see Fig. 4C. To this end, we vary the normalized cortical tension *γ*_*n*_, cortical viscosity *κ* and contact friction *λ* = *ξ*4*πR*_0_*/*(*γ*_*M*_*τ*_*d*_), where *ξ* represents the cell-cell viscous friction constant (Pa s m^−1^). We observe that most cell divisions are highly robust to changes in cell mechanical properties. However, a high relative cortex tension *γ*_*n*_ was found to influence cell volume, especially for the smaller cells which are characterized by high curvatures, such as ABpl. From a comparison between the P0 → AB + P1 division and the unconfined setup from the previous section, the stabilizing effect of geometrical confinement from the egg becomes apparent, see scaling in Fig. 3C,D versus Fig. 4C. The effect of relative cortex viscosity *κ* was found to be negligible, further supporting the notion of increased stability due to geometrical confinement. Similarly, varying the normalized contact friction *λ* produced no significant effects on cell volume ratio. This latter observation is to be expected, as up to the 6 cell stage embryo, cell motility is low and no drastic cell-cell contact area changes occur.

## Discussion

In this work, we presented a computational mechanical model of cytokinesis, which explicitly represents the contractile furrow ring that drives ingression of a cortical shell under mitotic tension. We demonstrated that this model well describes the dynamics of furrow ingression, on the condition that the furrow ring is mechanically dominant compared to the cortex surface tension and effective viscosity. These results agree with the predictions of Turlier et al. who used a visco-active membrane theory of the cortex to provide a general framework for interpreting and characterizing constriction failure and furrow ring closure as a result of underlying furrow ring and cortex mechanics [22]. Furthermore, we extended the cytokinesis model to account for asymmetric cell division by varying the position and angle of the mitotic spindle complex. In case division mechanics are dominant over cortex mechanics, the model reproduces a simple geometric relationship between spindle offset and daughter volume fraction. When confined between parallel plates, daughter cell asymmetry increases when the spindle orientation is parallel to the plate and decreases when it is perpendicular to the plate. For uniaxial compression, these differences in sensitivity can be traced back to a change in the characteristic length in the direction perpendicular to the division plane. For non-trivial compressed states, e.g., cells during *C. elegans* embryogenesis, similar effects occur where the division direction and characteristic length heavily influence the robustness of the system to any perturbation in spindle positioning. For *C. elegans* specifically, this partially explains the observed changes in volume ratio for ABa and ABp daughter cells.

We demonstrated that that the model is able to recapitulate *C. elegans* embryogenesis up to the six cell stage, using spindle position and angle as model input. In the case of *C. elegans*, we conclude that spindle position alone predicts the observed volume fraction after cell division very well. This supports the proposition that during cytokinesis, the mechanical properties of the contractile ring are dominant with respect to cortical properties such as mitotic tension and viscosity. In contrast to our initial hypothesis, the model predicts that differences in volume fraction between compressed and uncompressed embryos cannot be attributed solely to the effect of confinement on modeled cytokinesis mechanics alone. Rather, our result suggest that embryo compression also influences the positioning or angle of the spindle apparatus, an effect that is not accounted for by our model but may be expected based on cortical flows [10, 45]. Indeed, in a reverse fitting procedure, we estimated spindle positions based on the experimentally observed volume of cells, and predict a difference in spindle position between compressed and uncompressed embryos. The introduction of a non-zero bulk rigidity stabilizes cell divisions in the viscous cell model by preventing bubble-instabilities. Moreover, we predict that additional stability is provided by the eggshell. Indeed, the first division (P0) simulations are highly stable, even at greatly elevated mitotic surface tension. This observation highlights a possible additional role of the eggshell — or similar structures in higher organisms — during the cleavage phase of development. It does not only provide protection against outside chemical and mechanical stress, but also provides supplementary stability for asymmetric divisions.

We did not investigate the effects of cell-cell adhesion, cell-egg adhesion and embryo-egg volume ratio, as Giammona et al. have already covered these topics [28]. Finally, we also did not investigate the important effect of confinement, cell-cell connectivity and signaling on spindle offset *α* and skew angle *ϕ* [10, 45]. Incorporating the underlying mechanisms that mechanically regulate these structures will prove crucial for expanding this model towards fully predictive simulations of development. Nonetheless, this work shows the feasibility for the DCM framework to simulate developing systems such as early *C. elegans* morphogenesis, while allowing for detailed comparison to experimental observations.

## Models

We approximate the mechanical behavior of the cell cortex as a viscous shell, represented by a triangulated surface mesh where nodal positions act as the relevant degrees of freedom, see Fig. 2A. The cortical shell is parameterized by a thickness *t*, surface tension *γ* and viscosity *η*. Cell volume is conserved using an apparent bulk modulus *K*′. For simulation of multicellular *C. elegans* development, adhesive cell-cell and cell-egg contact with adhesion energy *w* (assumed ≪ *γ*) and wet friction constant *ξ* is implemented [32–34]. A full mechanical description of the model can be found in S1.

### Computational model of the furrow ring

Structurally, the furrow ring is composed of fixed length sub-units, in which tension is generated due to depolymerization of sub-units in the presence of end-tracking cross-linkers [17, 46]. To model this process, the furrow ring is represented as an elastic ring in the mother cell cortex, consisting of sub-units with a given initial length. Depolymerization is modeled by decreasing the resting length of the elastic sub-units, locally generating tensile stress in the cortex [17]. We do not model furrow-microtubule interactions, but introduce an effective sub-unit depolymerization velocity as

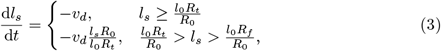

with base depolymerization velocity 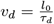, given initial sub-unit lengths *l*_0_ = ±600 nm and base ring closure time *τ*_*d*_ [17, 36]. Integrating eq. 3 yields eq. 1, as fit on experimental data by Carvalho et al., when fixing *R*_*t*_ = 3.6 μm and *R*_*f*_ = 0.8 μm. Using a linear elastic material model for the furrow ring, the in-plane elastic energy is expressed as

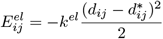

where *d*_*ij*_ = ‖***x***_*i*_ − ***x***_*j*_‖ and 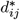 represent the current and resting distance between nodes *ij* and where *k*^*el*^ is the spring stiffness. The spring force between connected nodes is then given by

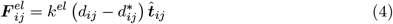

with direction

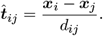

Consequently, the nodal spring forces are 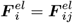 and 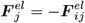. As nodal positions do not necessarily align with sub-unit endpoints, the furrow ring is approximated by a closed loop consisting of node-node connections closest to the division plane. For a given mesh refinement, this means that the selected node-node edges can be either larger or smaller than *l*_0_. To address this scaling issue, the effective node-node depolymerization velocity

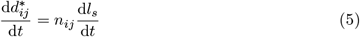

is determined for each unique connection, with scale factor

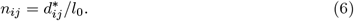

The spring force 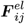 is added to the equation of motion presented in S1. Additionally, the local viscosity is increased accordingly, see S1, eq. 5. The local force balance induces cortex deformation, resulting in furrow ingression, see Fig. 1. Cell abscission is triggered when the furrow perimeter reaches a set threshold value, 2*πR*_*f*_. During this phase, furrow ring components are disassembled and the mother cell is split into two new entities, where the daughter cells inherit the mechanical properties of the mother cell. Simulated constriction velocities were estimated based on numerical differentiation of the ring perimeter with a step size of 10 s, while the total cytokinesis duration was estimated by taking the difference between cell abscission and cytokinesis initiation timings.

### Stabilization of polar instability in dividing cells

Asymmetric division of a viscous tensile shell is mechanically inherently unstable due to polar contractility, resulting in the bubble-instability. As we do not account for microtubule-cortex interactions, we use the approach proposed Sedzinski et al., where a linear bulk elastic resistance *K*^*div*^ is introduced to account for these interactions, assuring cell shape stabilization and avoiding daughter-half collapse during cytokinesis [31]. This stabilization model assumes cytosolic flow between daughter halves to be tightly regulated so that daughter halves can be treated as individual volumes. Using a proportional pressure controller

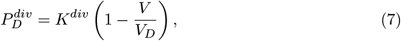

the reference volume *V*_*D*_ of daughter-half *D*, based on the division plane and mother cell polarization, can be enforced. 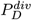 is integrated over the nodal Voronoi area *S*_*i*_ and added to equation of motion, S1, eq. 3. The net reaction force due to the pressure difference between daughter halves,

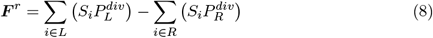

is applied as a reaction force to the furrow ring. This ensures that a proper force balance is maintained at the global (cell) level.

## Supporting information

**S1 Viscous deformable cell model**. Complete overview of the deformable cell model used in this work.

**S2 Supplementary information**. Extra data and figures.

**S3 Simulation visualizations**. Visualization of simulation output for the different setups used in this work.

## Author contributions

M. C. and B. S. conceived the project. M. C. and B. S. designed and conducted the simulations, M. C. and B. S. performed data analysis. W.T and R.J. contributed ideas and data for the application to *C. elegans* embryogenesis. M. C., W. T., R. J. and B. S. wrote the manuscript.

## Acknowledgments

The authors thank Prof. Matthias Weiss from the Experimental Physics I group, University of Bayreuth (Germany) for providing the experimental spindle offset measures.

This work is part of Prometheus, the KU Leuven R&D Division for Skeletal Tissue Engineering. M. C. acknowledges support form the Research Foundation Flanders (FWO), grant 1S46817N. W. T. acknowledges support form the Research Foundation Flanders (FWO), grant 11I2921N. R.J acknowledges support from the Research Foundation Flanders (FWO), grant G055017N. B. S. acknowledges support from the Research Foundation Flanders (FWO) grant 12Z6118N, and KU Leuven internal funding C14/18/055.

## Notes

### Competing Interest Statement

The authors have declared no competing interest.

